# Transcriptomic profiling of fibropapillomatosis in green sea turtles (*Chelonia mydas*) from South Texas

**DOI:** 10.1101/2020.10.29.360834

**Authors:** Nicholas B. Blackburn, Ana Cristina Leandro, Nina Nahvi, Mariana A. Devlin, Marcelo Leandro, Ignacio Martinez Escobedo, Juan M. Peralta, Jeff George, Brian A. Stacy, Thomas W. deMaar, John Blangero, Megan Keniry, Joanne E Curran

**Author notes:** **Correspondence to:** Nicholas B. Blackburn. Email contact and.

## Abstract

Sea turtle fibropapillomatosis (FP) is a tumor promoting disease that is one of several threats globally to endangered sea turtle populations. The prevalence of FP is highest in green sea turtle (*Chelonia mydas*) populations, and historically has shown considerable temporal growth. FP tumors can significantly affect the ability of turtles to forage for food and avoid predation and can grow to debilitating sizes. In the current study, based in South Texas, we have applied transcriptome sequencing to FP tumors and healthy control tissue to study the gene expression profiles of FP. By identifying differentially expressed turtle genes in FP, and matching these genes to their closest human ortholog we draw on the wealth of human based knowledge, specifically human cancer, to identify new insights into the biology of sea turtle FP. We show that several genes aberrantly expressed in FP tumors have known tumor promoting biology in humans, including *CTHRC1* and *NLRC5*, and provide support that disruption of the Wnt signaling pathway is a feature of FP. Further, we profiled the expression of current targets of immune checkpoint inhibitors from human oncology in FP tumors and identified potential candidates for future studies.

## INTRODUCTION

Sea turtle fibropapillomatosis (FP) is a tumor promoting disease that occurs globally in all seven species of sea turtles. The species known for having the highest FP prevalence is the green sea turtle (*Chelonia mydas*).^1^ FP was first reported over 80 years ago in captured green sea turtles from the Florida Keys, USA.^2^ Despite sustained and/or increased detection in Florida^3,4^, Hawaii^5–7^ and other areas globally^8,9^, FP was not detected in Texas, USA until 2010.^10^ Since then the prevalence of this disease in Texas has increased dramatically. From 2010 to 2015 the yearly prevalence of FP in Texas among stranded turtles was a maximum of 5% each year.^11^ After 2015, the prevalence of FP in Texas grew substantially to over 35% in green sea turtles in 2018.^11^

The most visibly prominent feature of FP is the appearance of external cutaneous tumors on affected turtles (Figure 1).^12–15^ Ocular, oral and internal tumors are also a disease component, affecting the visceral organs including the lungs, kidneys, and also bone.^16^ Juvenile sea turtles are recognized as the most affected age category, but FP has also been reported in adult nesting olive ridley turtles (*Lepidochelys olivacea*) in Costa Rica^17^ indicating that this disease is not limited to a life stage. Tumors may spontaneously regress, or increase in size and/or number to the point of debilitation.^18^ The primary impact of FP is the decrease in ability of sea turtles to forage for food, swim, and to avoid predation, thus affecting the overall survival of affected turtles.^16^ As green sea turtles are recognized by the International Union for the Conservation of Nature and Natural Resources as threatened with extinction^19^ health problems like FP that affect remaining populations are of major concern to conservation efforts.

**Figure 1.**
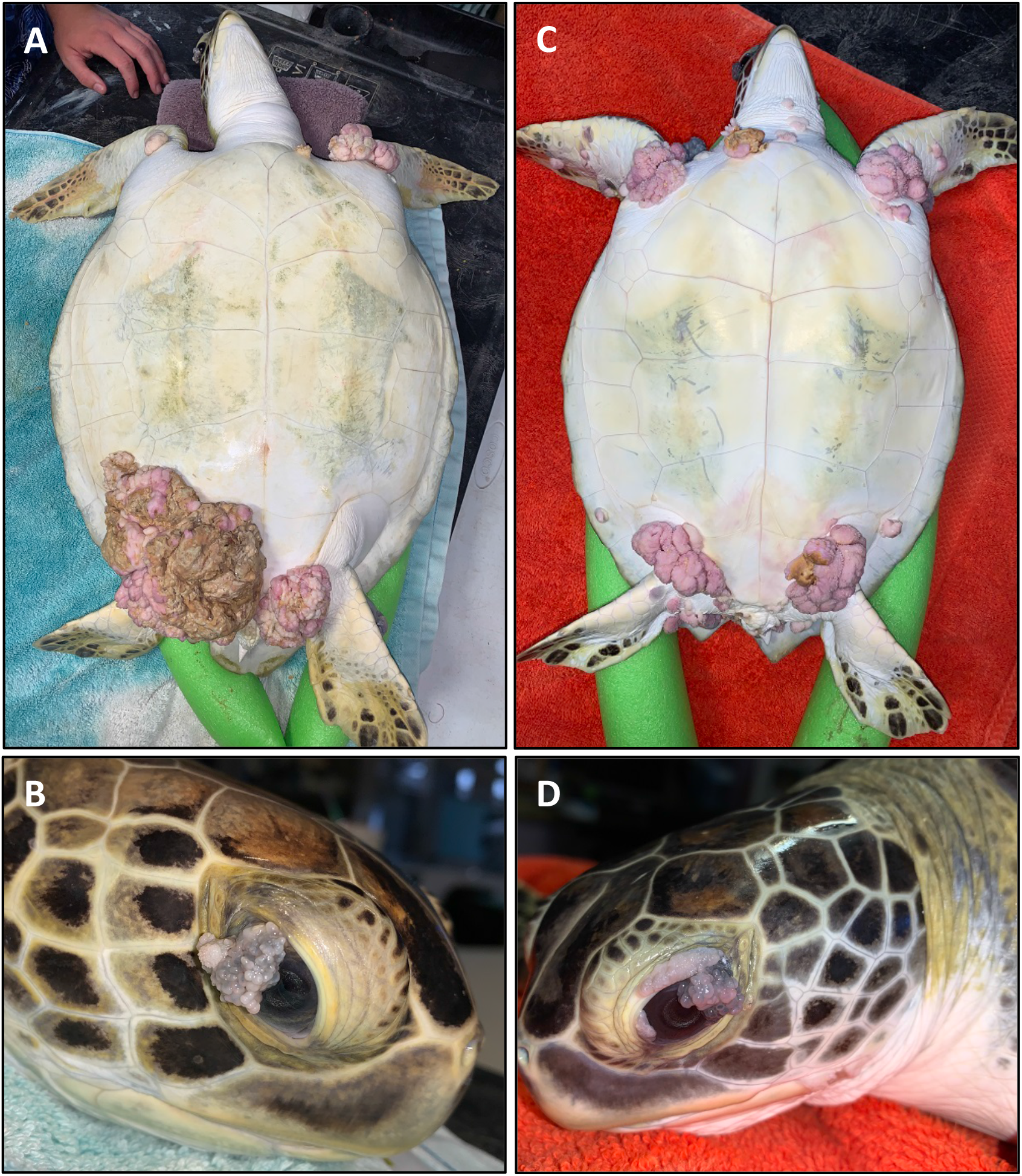
Two fibropapillomatosis affected juvenile green sea turtles (*Chelonia mydas*) undergoing rehabilitation at Sea Turtle Inc., South Padre Island, Texas, USA. Turtle 1 (A, B) and turtle 2 (C, D) both display prominent external tumors on their ventral side (A, C) and also ocular tumors (B, D).

The primary cause of FP is unknown. Several lines of evidence point to a herpesvirus (chelonid alphaherpesvirus 5 [ChHV5]) as a potential causal agent of FP^4,20–23^, which is reminiscent of human cancers with known viral origins including Epstein-Barr virus infection in Burkitt’s lymphoma^24^, human herpesvirus-8 infection in Karposi sarcoma^25^ and human papillomavirus infection in cervical cancer^26^. While mounting evidence implicates ChHV5 as at least a contributing risk factor for FP development, the presence of the virus in unaffected turtles^27^, together with the as yet unfulfilled Koch’s postulates indicates the need for more work to strengthen the causal association. Further there is a strong suggestion that it is ChHV5 together with other environmental and anthropogenic pressures that has contributed to the growth of FP prevalence (reviewed in Jones et al. 2016^1^).

For sea turtles debilitated by FP that are admitted to rehabilitation facilities for treatment, the current standard of care is preoperative screening for internal tumors, which carry poor prognosis, followed by surgical excision.^28^ A recent report from Page-Karjian et al. shows that postoperative regrowth is seen in 50% of treated turtles and that the rehabilitation survival rate of FP affected turtles is low (25%).^15^ Given the significant advancements in human precision oncology and the potential applications in wildlife medicine^29–31^ it seems reasonable to expect that if the understanding of the biological features of FP increases this may result in additional therapeutic approaches to the treatment of this disease. In turn this may increase survival rates, reduce tumor reoccurrence and minimize rehabilitation time. Moreover, insight into pathogenesis of tumor development may inform hypotheses related to disease etiology and expression.

In 2018 Duffy et al. reported the first application of human precision oncology techniques to FP.^32^ The study undertook a molecular characterization of FP tumors in two juvenile green sea turtles from Florida, performing RNA-sequencing of seven tumor samples and three matched skin control samples. This was the first effort to characterize the genetic disruptions in FP and was a significant advancement in the molecular understanding of this disease. More recent work from this group has explored the shared and distinct genomic and transcriptomic profiles of internal and external FP tumors.^33^

Here we continue the efforts to understand the genetic perturbations of FP tumors through transcriptome sequencing of healthy and tumor tissue from stranded green sea turtles undergoing rehabilitation at Sea Turtle Inc. on South Padre Island, Texas, USA. We aimed to study FP in a geographically distinct population of green sea turtles, hypothesizing that both similarities and differences will emerge between the transcriptomic profiles of FP tumors in Texas and Florida. By drawing upon the wealth of biological knowledge established in humans this study aimed to further characterize FP to understand the biological pathways disrupted in this disease and to identify potential therapeutic avenues for future investigation.

## RESULTS

### Patient characteristics

35 tissue samples were collected from nine green sea turtles undergoing rehabilitation at Sea Turtle Inc. after stranding at South Padre Island, Texas. As summarized in Table 1, 26 FP tumors and three healthy tissue biopsies were collected from three turtles with FP. Healthy tissue biopsies were also obtained from six turtles with no visible evidence of FP.

**Table 1:** Turtle characteristics.

| Table 1: Turtle characteristics |  |  |  |  |  |  |  |  |  |  |  |
| --- | --- | --- | --- | --- | --- | --- | --- | --- | --- | --- | --- |
| ID | Turtle | FP | Stranding date | Stranding reason | SCL N-T<sup>a</sup> (cm) | Weight (kg) | Body condition score<sup>b</sup> | FP score | Number of FP surgeries | Tumor samples | Healthy tissue samples |
| FP01 | Reveille | Yes | 1/30/18 | Head injury and eye injury | 29.4 | 6.05 | 3.5 | 1 | 5 | 3 | 1 |
| FP02 | Frostbite | Yes | 1/23/18 | Cold stun | 40.9 | 9.95 | 3 | 1 | 3 | 4 | 1 |
| FP03 | Beach Bum | Yes | 1/30/18 | Lethargic in tide line | 41.7 | 8.6 | 2 | 2 | 2 | 19 | 1 |
| FP07 | Robocop | No | 1/2/18 | Cold stun | 55.7 | 20.6 | 2.5 | NA | NA | NA | 1 |
| FP08 | Rocko | No | 4/2/18 | Hooked | 35.6 | 6.1 | 3 | NA | NA | NA | 1 |
| FP09 | Gobbler | No | 11/23/17 | Boat strike | 32.2 | 6.2 | 1.5 | NA | NA | NA | 1 |
| FP10 | Captain Cloud | No | 3/26/18 | Hooked | 30.1 | 3.25 | 3 | NA | NA | NA | 1 |
| FP11 | Sapphire | No | 2/8/18 | Cold stun | 27.4 | 3.1 | 3 | NA | NA | NA | 1 |
| FP12 | Gilligan | No | 1/16/18 | Cold stun | 27.6 | 3.5 | 4 | NA | NA | NA | 1 |

<sup>a</sup>Straight carapace length, notch to tip. <sup>b</sup>Body condition score is measured on a 1-5 scale, where 1 is considered extremely emaciated and 5 is considered obese.

The most common cause of stranding among the sampled sea turtles was cold stunning (4/9 turtles). The remaining turtles stranded due to trauma from injury (2/9), fishing hook ingestion / entanglement (2/9) and lethargy (1/9). All turtles were juveniles with straight carapace lengths (nuchal notch to caudal extent of the carapace) and weight ranging from 27.4 cm to 55.7 cm (μ=35.6 cm) and 3.1 kg to 20.6 kg (μ=7.5kg) respectively. For the three turtles that had FP tumors at collection, using the NOAA tumor burden scale of 1-3 (where 1 is least affected and 3 is most affected) two turtles were classified as category 1 and one turtle was category 2.^34^ All FP affected turtles underwent multiple tumor excisions by CO_2_ laser, with FP03 requiring two surgeries due to tumor load and both FP01 and FP02 requiring multiple surgeries due to tumor regrowth. All turtles were rehabilitated successfully at Sea Turtle Inc. and released.

General features observed from histopathology profiling of tumors collected and sequenced in this study confirmed samples as FP and identified common concurrent features including mild to moderate lymphocytic infiltration and ulceration with secondary bacterial infection in most tumors.

### Differential gene expression analysis of healthy vs tumor tissue

Our analysis of gene expression compared eight healthy control skin samples to 25 tumor samples across eight turtles total. Two samples, both tumors, from the originally sequenced 35 were excluded during quality control. Of the 21,432 transcripts annotated in the CheMyd 1.0 reference genome, 17,698 (82.6%) had sufficiently large enough counts in this sample set to be retained for statistical analysis. Using DESeq2, the design of this analysis controlled for 2 factors of unwanted variation calculated with RUVseq as well as the individual turtle (to account for differences in the number of tumor samples obtained from each FP turtle). As expected, principal components analysis of global gene expression profiles for each sample clustered distinctly healthy control skin samples separately from FP tumors (Figure 2), separating the two sample types in the first principal component which explained 24.62% of sample variation.

**Figure 2.**
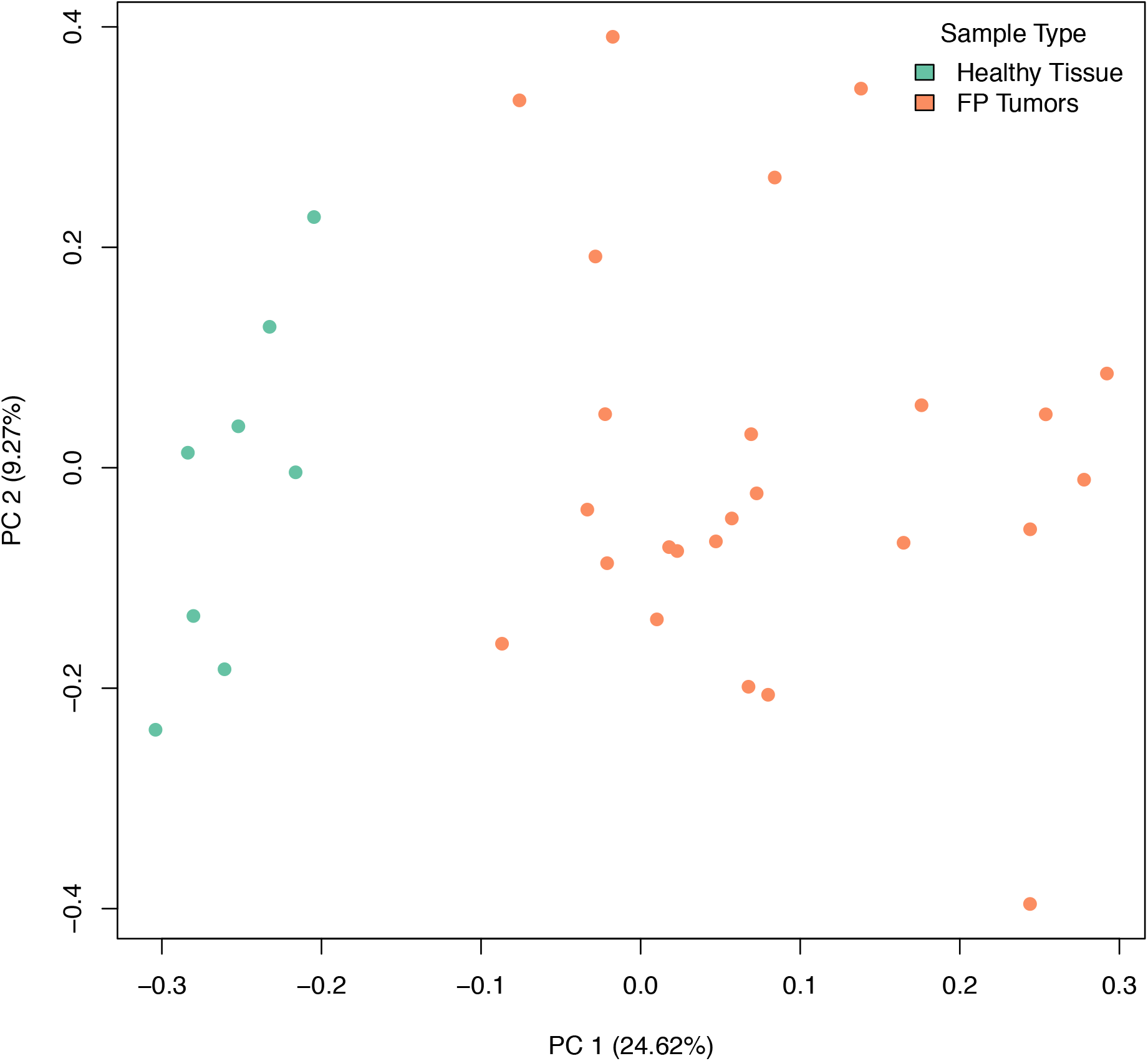
Principal components analysis of gene expression in healthy (green) and tumor (orange) samples, incorporating 2 factors of unwanted variation calculated using RUVSeq. PC 1 separates healthy from tumor samples and explains 24.62% of the variation in gene expression across samples.

For differential gene expression analysis, a log fold change threshold of 1 and an alpha of 0.05 were chosen. This analysis identified 283 transcripts that were upregulated, and 90 transcripts that were downregulated (Figure 3). The 10 most significantly upregulated, and 10 most significantly downregulated genes are shown in Figure 4 and described in Table 2. All gene expression differences are summarized in supplementary table 1.

**Table 2.** Top 10 upregulated and downregulated differentially expressed genes.

|  |  |  |  |  |
| --- | --- | --- | --- | --- |
| CheMyd_1.0<br>Gene Symbol | Human Ortholog<br>Gene Symbol | log2FoldChange | lfcSE | padj |
| upregulated |  |  |  |  |
| *LOC102930327* | *MR1* | 2.92 | 0.17 | $1.22 \times 10^{-25}$ |
| *CTHRC1* | *CTHRC1* | 6.60 | 0.53 | $6.06 \times 10^{-22}$ |
| *TBX20* | *TBX20* | 21.75 | 2.15 | $2.63 \times 10^{-18}$ |
| *LOC114021482* | *RNF213* | 3.47 | 0.26 | $1.69 \times 10^{-17}$ |
| *LOC102948206* | N/A | 16.86 | 1.89 | $1.29 \times 10^{-13}$ |
| *NLRC5* | *NLRC5* | 3.39 | 0.30 | $3.71 \times 10^{-12}$ |
| *LOC114018352* | *UBB* | 4.26 | 0.42 | $8.19 \times 10^{-12}$ |
| *CD4* | *CD4* | 4.49 | 0.46 | $3.68 \times 10^{-11}$ |
| *LOC102931356* | *CASP8* | 3.50 | 0.33 | $9.16 \times 10^{-11}$ |
| *TMSB4X* | *TMSB4X* | 2.42 | 0.20 | $1.03 \times 10^{-9}$ |
| downregulated |  |  |  |  |
| *TBX4* | *TBX4* | -7.35 | 0.71 | $1.00 \times 10^{-15}$ |
| *CCER2* | *CCER2* | -7.76 | 0.87 | $1.99 \times 10^{-11}$ |
| *LOC114020549* | *BTAF1* | -5.63 | 0.64 | $7.41 \times 10^{-10}$ |
| *LGR5* | *LGR5* | -4.56 | 0.50 | $1.03 \times 10^{-9}$ |
| *LOC114020545* | *PLIN2* | -2.27 | 0.22 | $8.27 \times 10^{-6}$ |
| *LOC114021518* | N/A | -4.11 | 0.56 | $1.08 \times 10^{-5}$ |
| *LOC114021212* | N/A | -4.21 | 0.58 | $1.13 \times 10^{-5}$ |
| *ENPP6* | *ENPP6* | -3.91 | 0.53 | $1.66 \times 10^{-5}$ |
| *LOC102936807* | *KRT73* | -9.18 | 1.54 | $3.58 \times 10^{-5}$ |
| *LOC114019042* | N/A | -5.98 | 0.94 | $3.60 \times 10^{-5}$ |

**Figure 3.**
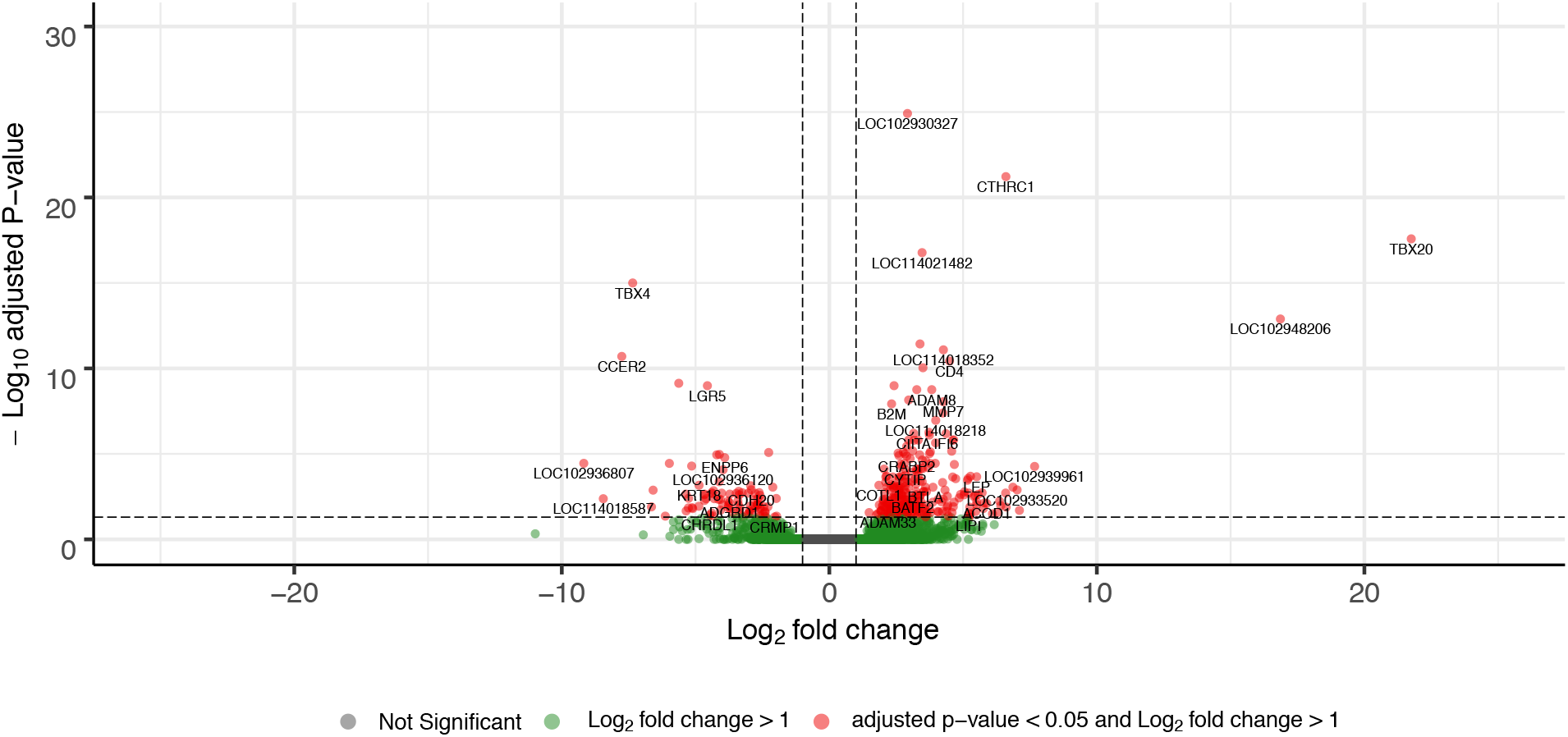
Volcano plot of differentially expressed genes in FP tumor samples compared to healthy control skin. The x-axis plots the log_2_ fold change from healthy control skin to tumor samples, while the y-axis plots the −log_10_ adjusted P-value resulting from the differential expression analysis.

**Figure 4.**
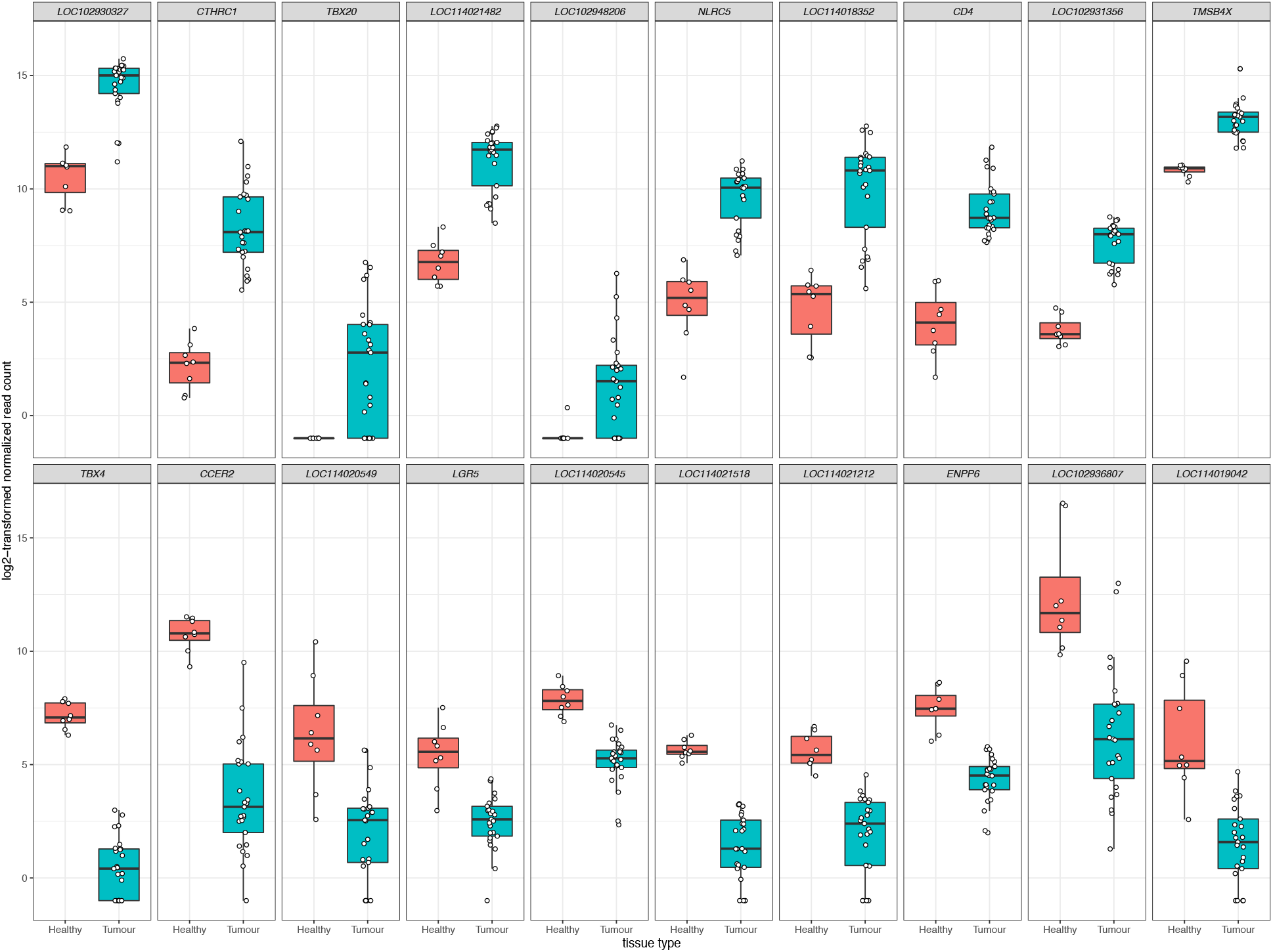
The top 10 upregulated (upper panels) and downregulated (lower panels) differentially expressed genes in FP tumors. The x-axis in each plot panel separates the box plots of tissue samples into the healthy control skin group and tumor group. The y-axis plots the log_2_-transformed normalized read count for each sample.

### Functional enrichment and pathway-based analysis of differentially expressed genes

To extend beyond individual differentially expressed genes and to examine the broader biological processes involved in FP tissue, turtle genes were mapped to their closest human gene ortholog. This then enabled analyses that draw upon well characterized biological networks from human based data to infer understandings in sea turtle FP.

Among the 17,698 sea turtle genes that underwent differential expression analysis between healthy tissue and tumors, 91.2% were successfully matched to a human ortholog through BLAST based analysis. For the 373 significantly differentially expressed genes, 327 (87.7%) were matched to 302 human orthologs.

GSEA using GO terms and KEGG pathways was conducted using clusterProfiler. For this analysis we relaxed the significance threshold and considered genes with an adjusted p-value < 0.1 as input on the basis that biological relevant expression changes may be reflected in this expanded list. This increased the number of genes to 459 of which 407 (88.7%) were matched to 374 human orthologs. For each duplicate ortholog the most significant differentially expressed gene value was used in downstream analyses.

From the GO based analysis, 465 nominally significant (p < 0.05) terms were identified as enriched within the 374 genes (supplementary table 2). The top ranked terms were primarily immune system related. The top 20 most significant GO biological process ontology terms are shown in Figure 5. When the genes overlapping between these GO terms are considered as an enrichment map (Figure 6), two functional modules emerge. One module specifically involves biological processes related to responses to bacteria and interferon-gamma and the second module is related to immune cells and immune processes.

**Figure 5.**
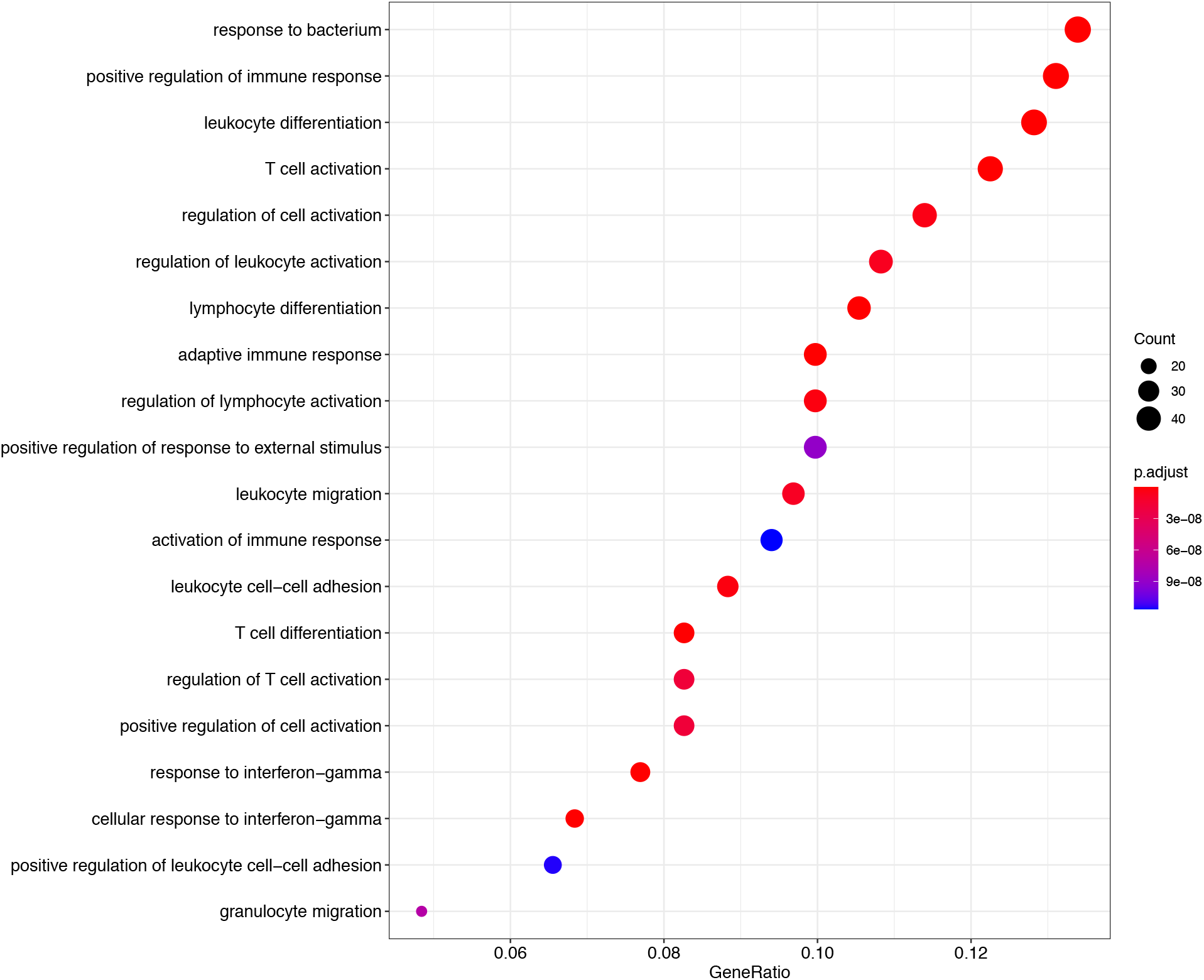
A dot plot of the top 20 most significant GO biological process ontology terms enriched among differentially expressed genes in FP. The x-axis shows the ratio of differentially expressed genes in each term relative to the total number of genes in that term. Dot size shows the number of differentially expressed genes in that term and color is the gene set enrichment analysis test statistic with Benjamini-Hochberg adjustment for multiple testing.

**Figure 6.**
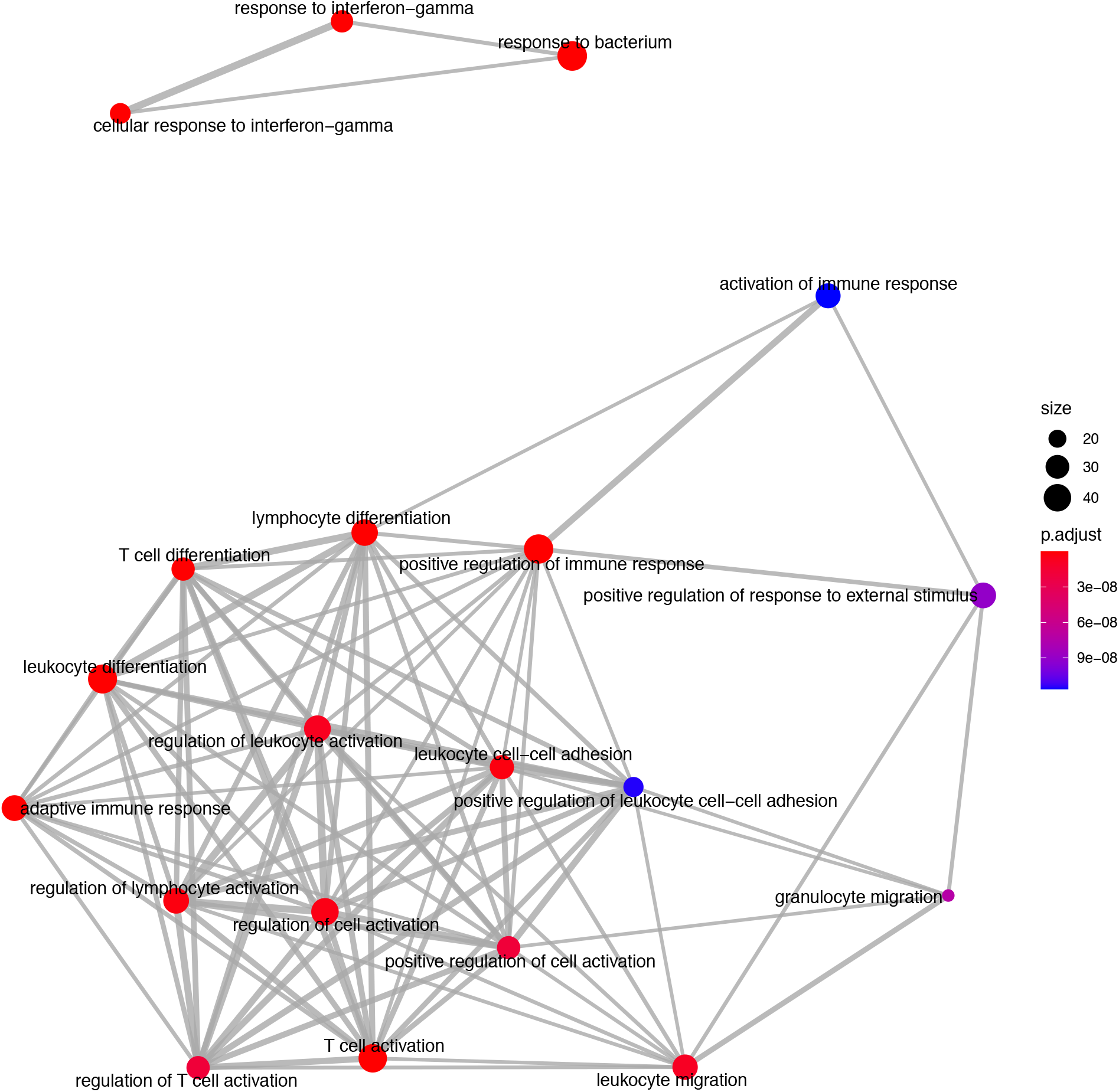
An enrichment map of genes overlapping the top 20 most significant GO biological process ontology terms enriched among differentially expressed genes in FP. The size of each node represents the number of genes differentially expressed in that ontology term, with thickness of edges between nodes representing the number of genes overlapping. Node color represents the gene set enrichment analysis test statistic with Benjamini-Hochberg adjustment for multiple testing.

From the KEGG pathway based analysis, 32 nominally significant pathways, shown in Figure 7 (supplementary table 3), were identified as over-represented within the differentially expressed genes. The most significantly enriched pathway was hsa04060, cytokine-cytokine receptor interaction, adjusted p = 8.72 × 10^−7^. Again, considering an enrichment map (Figure 8) between these pathways it is clear that immunological processes are driving the differential expression of genes between FP tumors and normal tissue.

**Figure 7.**
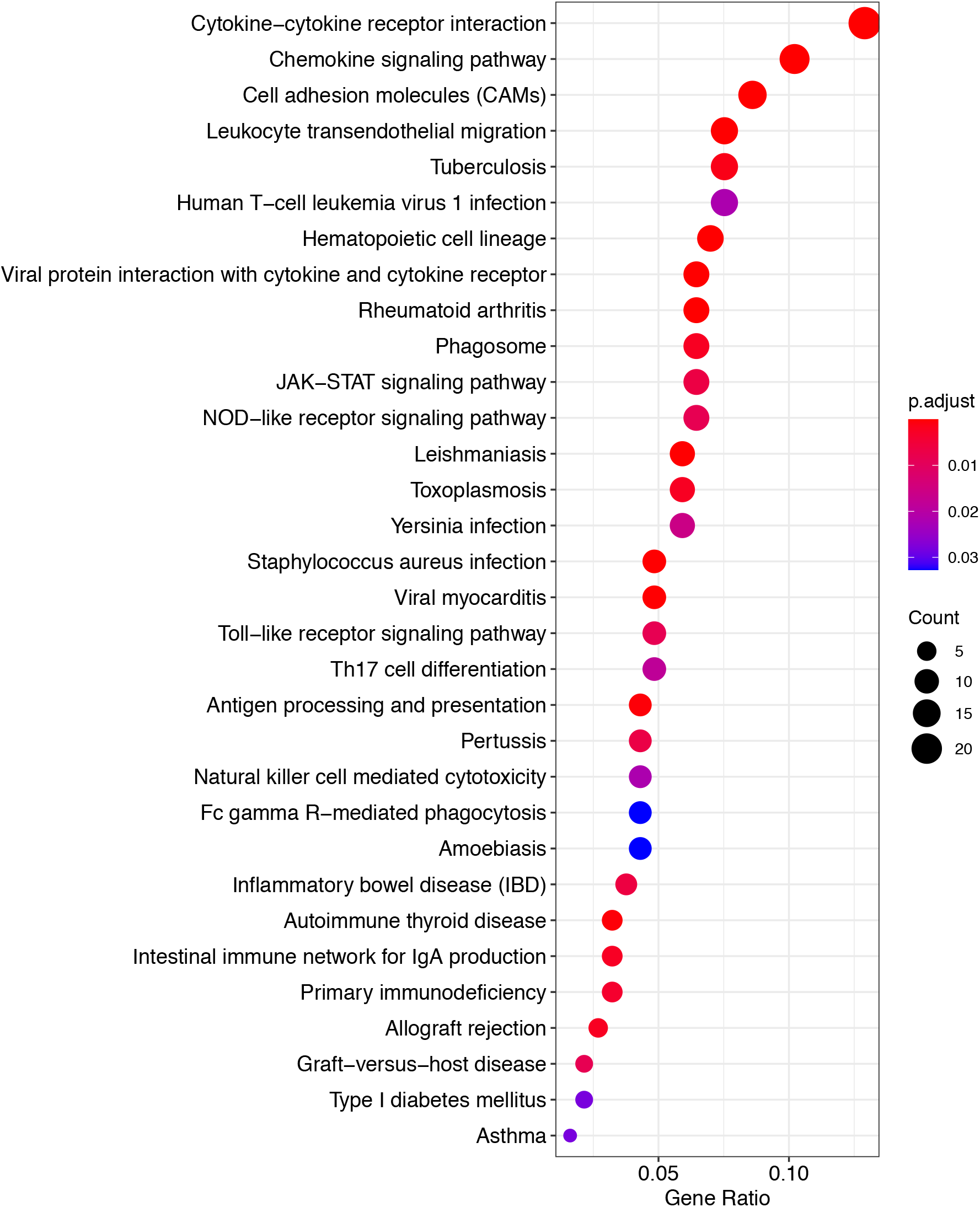
A dot plot of nominally significant KEGG pathways enriched among differentially expressed genes in FP. The x-axis shows the ratio of differentially expressed genes in each term relative to the total number of genes in that term. Dot size shows the number of differentially expressed genes in that term and color is the gene set enrichment analysis test statistic with Benjamini-Hochberg adjustment for multiple testing.

**Figure 8.**
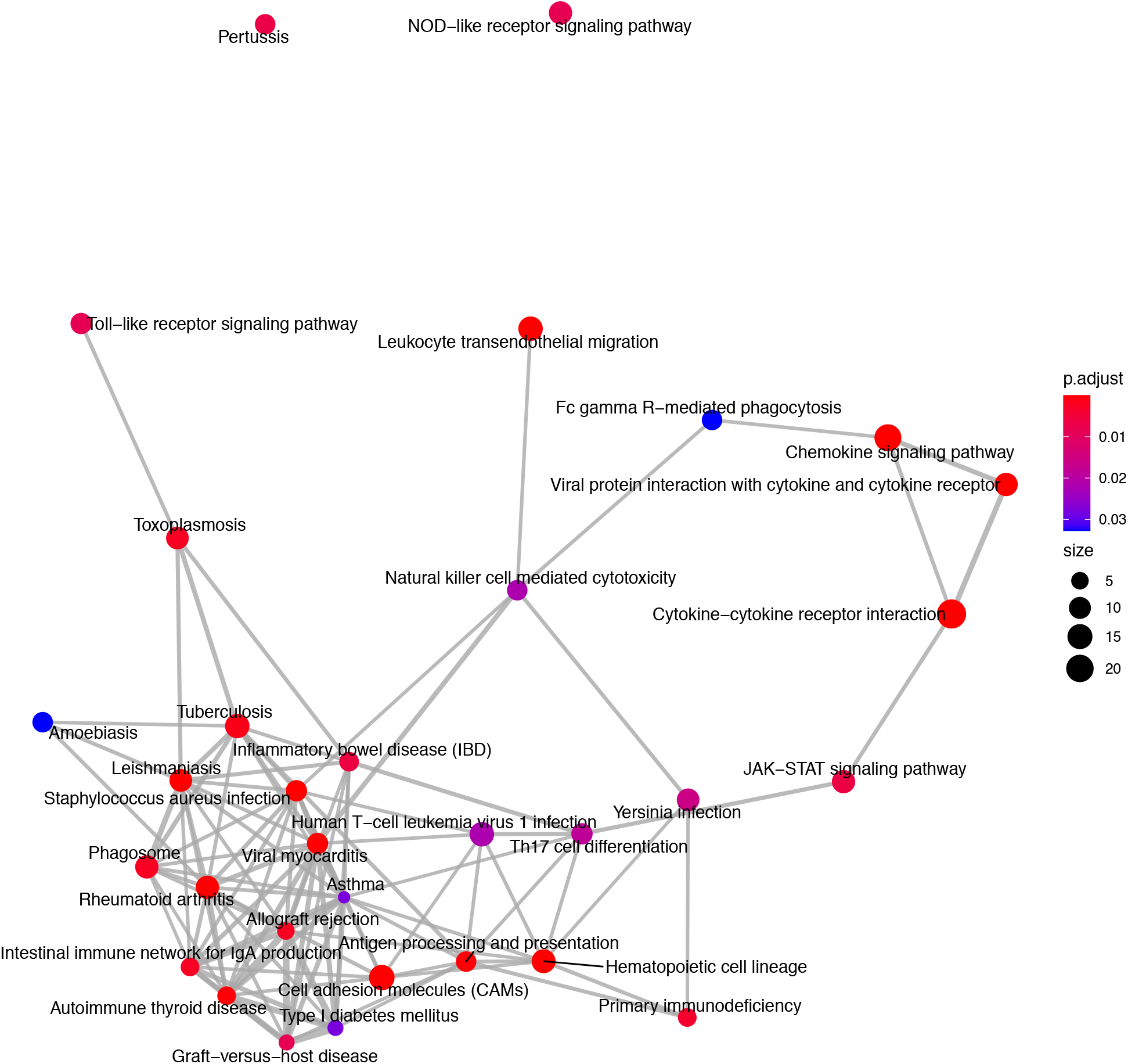
An enrichment map of genes overlapping nominally significant KEGG pathways enriched among differentially expressed genes in FP. The size of each node represents the number of genes differentially expressed in that KEGG pathway, with thickness of edges between nodes representing the number of genes overlapping. Node color represents the gene set enrichment analysis test statistic with Benjamini-Hochberg adjustment for multiple testing.

### Targeted analysis of immune checkpoint dysregulation in FP tumors

Inhibition of immune checkpoint molecules, including CTLA4 and PD1, are current therapeutic targets in human precision oncology. Given their high therapeutic potential we specifically assessed the gene expression of immune checkpoint molecule orthologs in this data including both established and emerging targets, as reviewed in De Sousa Linhares et al. and Waldman et al. and summarized in table 3.^35,36^ Of the 13 molecules targeted from the literature, 12 were present as orthologs in the sea turtle transcriptome based on protein sequence homology, with PD-L1, B7-H3 and CTLA-4 matching multiple sea turtle proteins. TIGIT was not detected. Three immune checkpoint molecules, (BTLA, PD-L2 and LAG3) were significantly upregulated in FP tumors (Figure 9, Table 4). The most significant being PD-L2 (*LOC102933512*), with a log2fold increase of 3.27, padj = 0.0001.

**Table 3.** Immune checkpoint molecules assessed.

| Table 3. Immune checkpoint molecules assessed |  |  |
| --- | --- | --- |
| Human gene symbol | Immune checkpoint molecule | Orthologous match with CheMyd_1.0 Gene Symbol/s |
| *BTLA* | BTLA | *BTLA* |
| *CD274* | PD-L1 | *CD274, LOC114018541* |
| *CD276* | B7-H3 | *LOC114022564, LOC114021416, LOC114022559, CD276, LOC102929593* |
| *CTLA4* | CTLA-4 | *LOC102947343, LOC114018946* |
| *HAVCR2* | TIM-3 | *LOC102941709* |
| *HHLA2* | B7-H7 | *HHLA2* |
| *LAG3* | LAG-3 | *LAG3* |
| *PDCD1* | PD-1 | *PDCD1* |
| *PDCD1LG2* | PD-L2 | *LOC102933512* |
| *TIGIT* | TIGIT | N/A |
| *TMIGD2* | CD28H | *LOC114018671* |
| *VSIR* | VISTA | *VSIR* |
| *VTCN1* | B7-H4 | *VTCN1* |

**Table 4.** Differential expression of immune checkpoint molecules in FP.

| Table 4. Differential expression of immune checkpoint molecules in FP |  |  |  |  |  |
| --- | --- | --- | --- | --- | --- |
| CheMyd_1.0 Gene Symbol | Human Ortholog Gene Symbol | Immune checkpoint molecule | log2FoldChange | lfcSE | padj |
| *LOC102933512* | *PDCD1LG2* | PD-L2 | 3.27 | 0.45 | 0.0001 |
| *BTLA* | *BTLA* | BTLA | 3.59 | 0.57 | 0.0009 |
| *LAG3* | *LAG3* | LAG-3 | 6.41 | 1.42 | 0.01 |
| *LOC102947343* | *CTLA4* | CTLA-4 | 4.39 | 1.11 | 0.09 |
| *LOC114022564* | *CD276* | B7-H3 | 1.99 | 0.36 | 0.19 |
| *LOC114021416* | *CD276* | B7-H3 | 3.74 | 1.00 | 0.20 |
| *CD274* | *CD274* | PD-L1 | 1.92 | 0.38 | 0.35 |
| *LOC102941709* | *HAVCR2* | TIM-3 | 1.81 | 0.35 | 0.46 |
| *LOC114022559* | *CD276* | B7-H3 | 3.14 | 1.10 | 0.83 |
| *LOC114018541* | *CD274* | PD-L1 | 0.49 | 0.71 | 1 |
| *CD276* | *CD276* | B7-H3 | 0.15 | 0.33 | 1 |
| *LOC102929593* | *CD276* | B7-H3 | 0.48 | 0.28 | 1 |
| *LOC114018946* | *CTLA4* | CTLA-4 | 2.05 | 0.87 | 1 |
| *HHLA2* | *HHLA2* | B7-H7 | 0.09 | 0.50 | 1 |
| *PDCD1* | *PDCD1* | PD-1 | 2.05 | 0.58 | 1 |
| *LOC114018671* | *TMIGD2* | CD28H | 1.16 | 0.43 | 1 |
| *VSIR* | *VSIR* | VISTA | 1.37 | 0.32 | 1 |
| *VTCN1* | *VTCN1* | B7-H4 | -0.75 | 0.61 | 1 |

**Figure 9.**
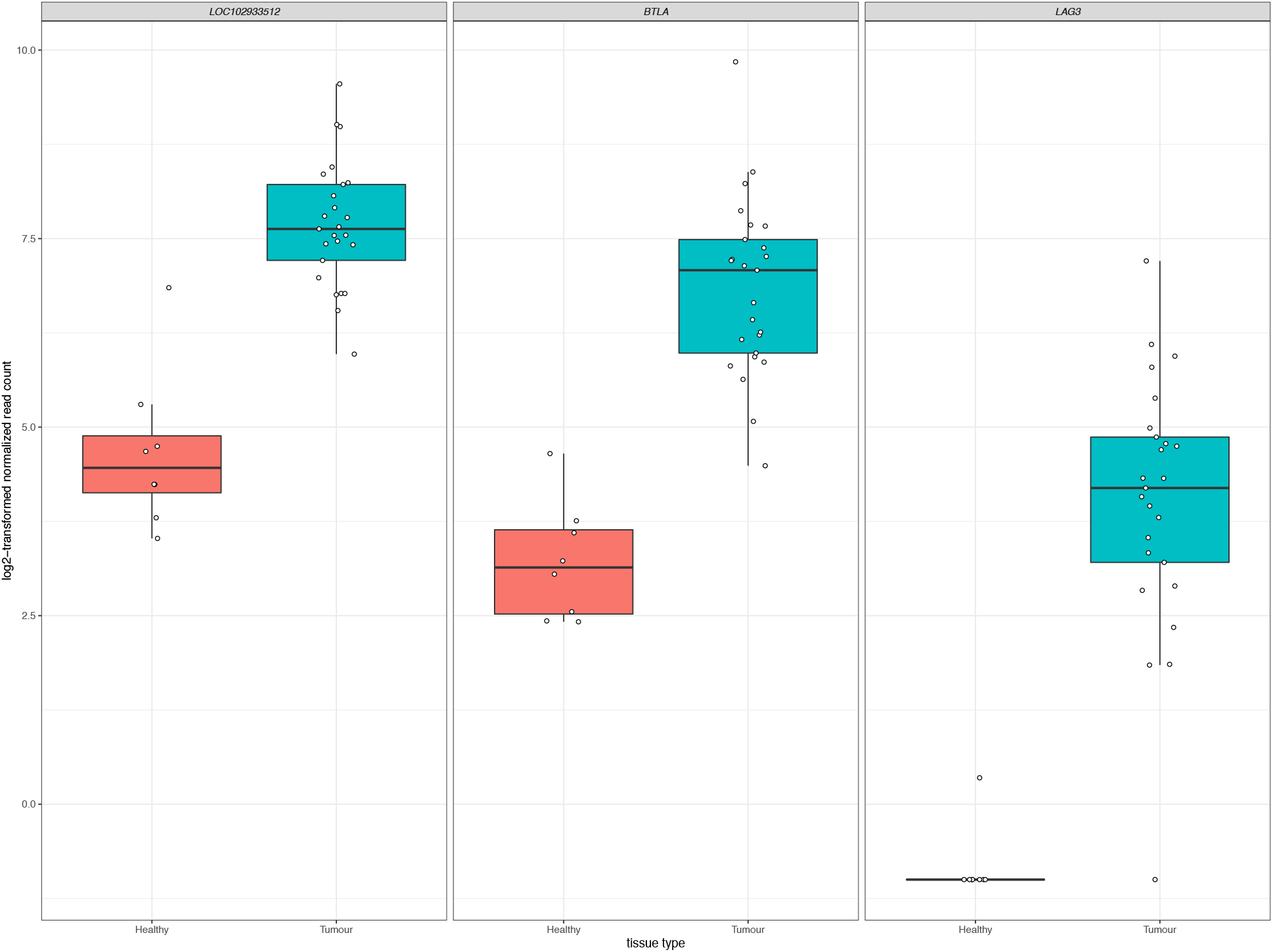
Significant upregulation of three immune checkpoint genes in FP. The x-axis in each plot panel separates the box plots of tissue samples into the healthy control skin group and tumor group. The y-axis plots the log_2_-transformed normalized read count for each sample.

### Similarities and differences between gene expression profiles of FP from sea turtles in Texas and Florida

Previously in the first reported transcriptomic profiling of FP, Duffy et al. sequenced 3 healthy control samples and 7 tumor samples from 2 green sea turtles in Florida^32^, assembled a *de novo* transcriptome and conducted a differential gene expression analysis. Here we compare our findings where possible with these earlier findings. Duffy et al. reported the top 10 up-regulated and top 10 down-regulated transcripts in their analysis of which 12 were annotated to a matching human ortholog. Table 5 compares the differential expression results for these 12 genes between the Duffy et al. turtles from Florida and the turtles in this current study. Of the 12 genes, only four (*FNDC1*, *S1PR3*, *NES*, *NTN3*) were statistically replicated in this study, however all showed the same direction of expression change.

**Table 5.**
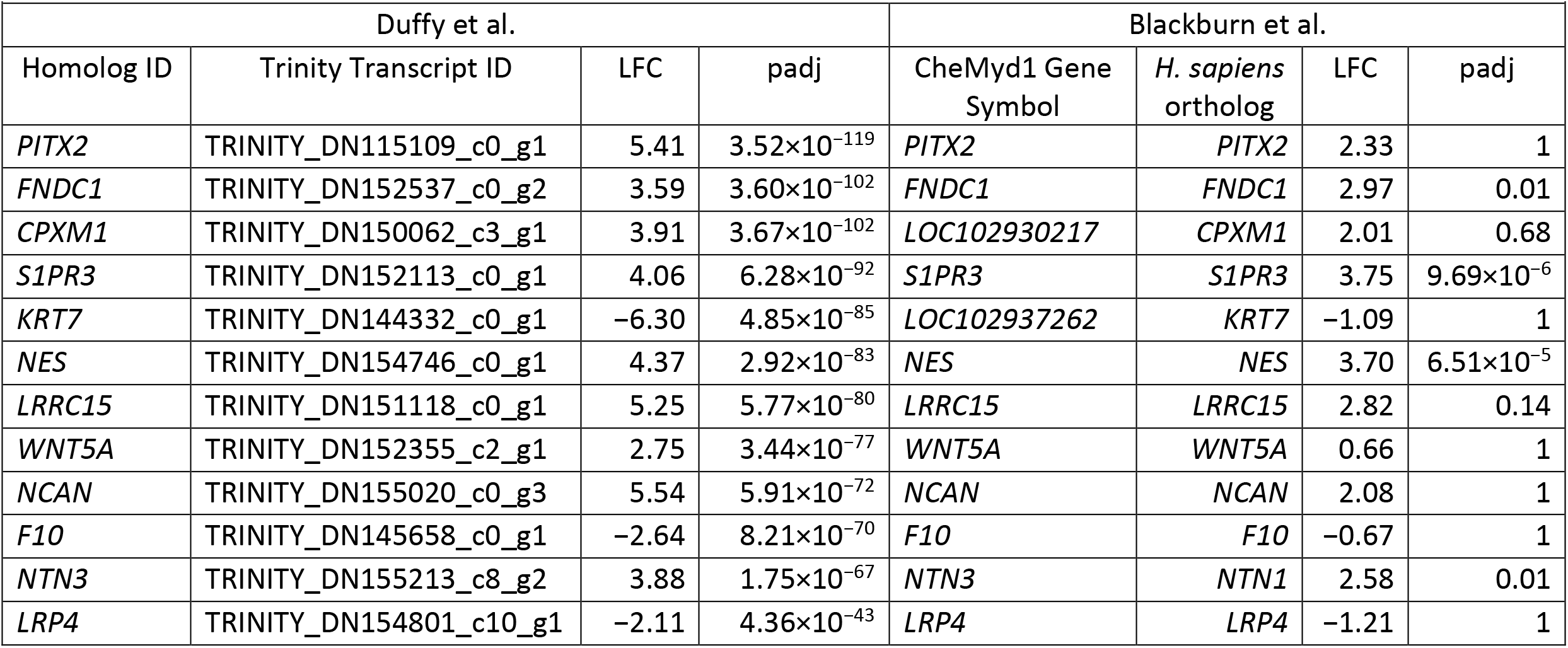
Comparison to previously identified top differentially expressed genes in FP from Duffy et al.

|  |  |  |  |  |  |  |  |
| --- | --- | --- | --- | --- | --- | --- | --- |
| Duffy et al. |  |  |  | Blackburn et al. |  |  |  |
| Homolog ID | Trinity Transcript ID | LFC | padj | CheMyd1 Gene Symbol | *H. sapiens* ortholog | LFC | padj |
| *PITX2* | TRINITY_DN115109_c0_g1 | 5.41 | $3.52 \times 10^{-119}$ | *PITX2* | *PITX2* | 2.33 | 1 |
| *FNDC1* | TRINITY_DN152537_c0_g2 | 3.59 | $3.60 \times 10^{-102}$ | *FNDC1* | *FNDC1* | 2.97 | 0.01 |
| *CPXM1* | TRINITY_DN150062_c3_g1 | 3.91 | $3.67 \times 10^{-102}$ | *LOC102930217* | *CPXM1* | 2.01 | 0.68 |
| *S1PR3* | TRINITY_DN152113_c0_g1 | 4.06 | $6.28 \times 10^{-92}$ | *S1PR3* | *S1PR3* | 3.75 | $9.69 \times 10^{-6}$ |
| *KRT7* | TRINITY_DN144332_c0_g1 | -6.30 | $4.85 \times 10^{-85}$ | *LOC102937262* | *KRT7* | -1.09 | 1 |
| *NES* | TRINITY_DN154746_c0_g1 | 4.37 | $2.92 \times 10^{-83}$ | *NES* | *NES* | 3.70 | $6.51 \times 10^{-5}$ |
| *LRRC15* | TRINITY_DN151118_c0_g1 | 5.25 | $5.77 \times 10^{-80}$ | *LRRC15* | *LRRC15* | 2.82 | 0.14 |
| *WNT5A* | TRINITY_DN152355_c2_g1 | 2.75 | $3.44 \times 10^{-77}$ | *WNT5A* | *WNT5A* | 0.66 | 1 |
| *NCAN* | TRINITY_DN155020_c0_g3 | 5.54 | $5.91 \times 10^{-72}$ | *NCAN* | *NCAN* | 2.08 | 1 |
| *F10* | TRINITY_DN145658_c0_g1 | -2.64 | $8.21 \times 10^{-70}$ | *F10* | *F10* | -0.67 | 1 |
| *NTN3* | TRINITY_DN155213_c8_g2 | 3.88 | $1.75 \times 10^{-67}$ | *NTN3* | *NTN1* | 2.58 | 0.01 |
| *LRP4* | TRINITY_DN154801_c10_g1 | -2.11 | $4.36 \times 10^{-43}$ | *LRP4* | *LRP4* | -1.21 | 1 |

Recognizing that the results from the two sample cohorts are derived from different sequencing and analytical pipelines we can compare the trend in overall log2fold changes in gene expression where possible. Duffy et al. report their top 300 up-regulated and top 300 down-regulated transcripts. Where a human gene annotation was assigned to a transcript, we have correspondingly matched that gene to our data, allowing for multiple matches where the same gene symbol is assigned to multiple transcripts. Figure 10A shows the direct comparison of log2fold changes in gene expression between 449 overlapping observations from Duffy et al. and this study. Two transcripts from the Duffy study, both corresponding to the *TBX20* gene were excluded as outliers as the *TBX20* gene had a log2fold change of 21.8 in our study, in comparison to 2.9 and 3.6 in the Duffy study. The overlap between studies was highly correlated (R = 0.76, p < 2.2×10^−16^), with a decrease in correlation to R = 0.68 with inclusion of *TBX20* data points. This high correlation was increased when considering only the subset of transcripts (N = 98) that were significantly differentially expressed in both studies (R = 0.86, p < 2.2×10^−16^) as seen in Figure 10B.

**Figure 10.**
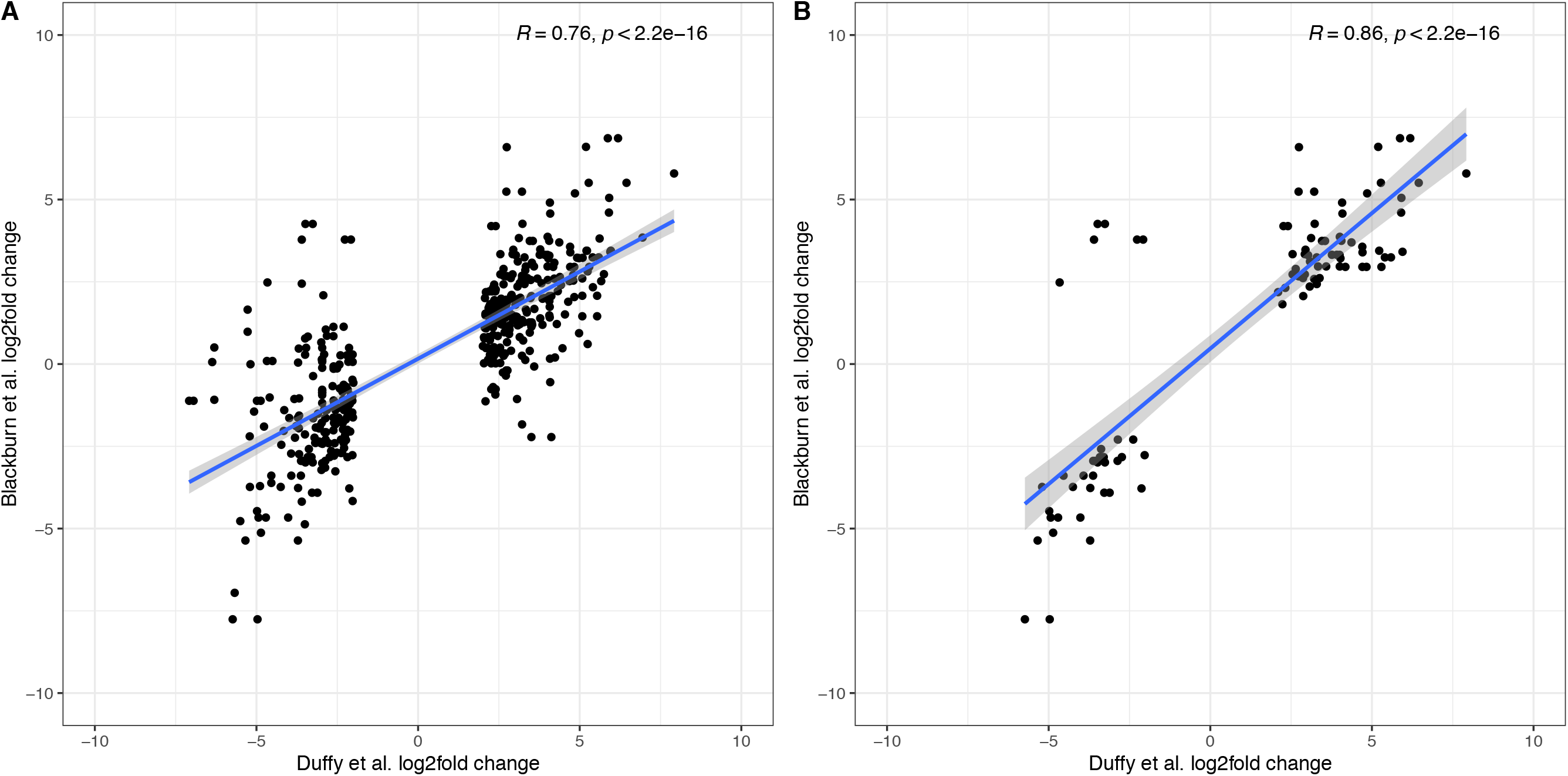
Comparison of differentially expressed genes reported in Duffy et al. to the current study. A) log2fold correlations between 449 overlapping transcripts between the two studies. B) log2fold correlations for the subset of overlapping transcripts that were differentially expressed in both studies.

### Detection of ChHV5 transcript expression in both tumor and normal tissue

We aligned the RNA sequence data from this study to the ChHV5 reference genome to identify sample level evidence of the presence of the ChHV5 virus. Of the 104 defined gene features in the ChHV5 genome, 83 were detected in this sample set. Total alignment counts across all 83 transcripts ranged from 0 to 483,216, with less than 10 counts in the majority of samples (22/33). Across the eight healthy tissue samples RNA molecules aligning to ChHV5 virus transcripts were detected in five, including all three healthy tissue samples from the FP affected turtles. In tumor samples alignment to virus transcripts was detected in 20/25 samples with two tumors from the same turtle (FP01) exhibiting high levels of RNA molecules aligning to ChHV5 transcripts. Five tumors, all from turtle FP03 had no evidence of ChHV5 transcript expression. We report with interest that the two tumors from turtle FP01 with the apparent highest virus levels exhibited a smooth, raised surface with altered pigmentation in comparison to surrounding normal tissue, in comparison to the more typical verrucous character of the other tumors in this study.

## DISCUSSION

The prevalence of FP in Texas is growing among juvenile green turtles inhabiting nearshore habitat. As a wildlife health problem threatening an endangered species there is considerable need for further biological insight into this disease. Here, we have conducted transcriptome sequencing to identify differential gene expression profiles between healthy skin samples and FP tumors. Our analysis has identified 373 significantly differentially expressed sea turtle transcripts between healthy and tumor tissue. Through protein sequence homology analysis, we identified, where possible, the best matching human ortholog for each transcript. This facilitated exploration of both known biology at the gene level, but also allowed GSEA to identify aberrantly affected biological processes and pathways.

At the gene level, among the most significantly up-regulated and down-regulated genes resulting from this analysis are several genes currently implicated as important in human oncology.

Expression of *LOC102930327* which was found to be orthologous with human *MR1* (major histocompatibility complex, class I-related) is upregulated in sea turtle FP tumors (log2FoldChange = 2.92, padj = 1.22×10^−25^). *MR1* has a known role in the innate immune system as a selector and/or restrictor of mucosal-associated invariant T (MAIT) cells, which have a role in microbial defense.^37–39^ Through antigen presentation by MR1, MAIT lymphocytes are activated by microbially-derived intermediates of riboflavin synthesis.^40^ MAIT cells are the dominant lymphocyte population in human skin and also have a role in promoting tissue repair and wound healing.^41^ As FP tumors have been observed to have bacterial infections previously^5^ as well as in the current study, we posit that the identified *MR1* upregulation in FP tumors is an antimicrobial response and while not directly related to FP tumorgenicity may contribute to a permissive tumor microenvironment through inflammation upregulation. It is possible that as FP tumors develop and tissue becomes compromised, facilitating bacterial infection, the infection in turn contributes to continued tumor growth.

Expression of *CTHRC1* (collagen triple helix repeat containing 1) is also upregulated in FP tumors (log2FoldChange = 6.60, padj = 6.06×10^−22^). In humans, *CTHRC1* encodes an extracellular matrix protein that was originally identified from an upregulation in models of arterial injury.^42^ Further research has led to the recognition of CTHRC1 as an oncogenic protein with a role in the promotion of tumor growth, invasion and metastasis in multiple malignancies. *CTHRC1* expression is increased in hepatocellular carcinoma^43,44^, melanoma^45^ and breast cancer^46^ and in several studies increased mRNA and protein expression in tumors was associated with poor disease prognosis and survival.^43,46–49^ Further, increased CTHRC1 has been linked to the upregulation of matrix metalloproteinases (MMPs) which have established roles in tumor progression and metastasis in humans.^50^ In heptoma cells^51^, and colorectal cancer cells^47^ increased CTHRC1 activated expression of MMP-9, which is known to contribute to extracellular matrix breakdown and tumor promotion.^50^ Increased expression of *MMP9* was associated with poorer outcomes in nasopharyngeal carcinoma patients^52^ and together with increased expression of *CTHCR1* and *MMP7*, higher expression of *MMP9* was associated with a lower survival rate after surgery in non-small cell lung cancer patients.^53^ This known connection between *CTHCR1* and *MMP7/MMP9* is reflected in the current study, with significant upregulation of *CTHRC1* in FP tumors accompanied by increased expression of *MMP7* (log2FoldChange = 4.26, padj = 8.87×10^−9^) and *MMP9* (log2FoldChange = 4.24, padj = 4.05×10^−8^) (supplementary table 1). Together this suggests that *CTHRC1* may be a key oncogenic driver in FP.

The most significantly downregulated gene expressed in FP tumors was *TBX4* (log2FoldChange = −7.35, padj = 1.00×10^−15^). Encoding T-box transcription factor 4, *TBX4*, belongs to a highly evolutionarily conserved family of transcription factors responsible for limb development with fundamental roles during embryogenesis.^54,55^ Given that *Tbx4* expression in chick embryos determines hind limb development^56,57^, and that the healthy control tissue samples used in the current project were obtained from punch biopsies of the hind flippers, it is possible that the higher expression of *TBX4* in healthy tissue in this study is due to tissue source. This finding warrants further investigation through gene expression comparisons of hind flipper tissue with non-hind flipper healthy skin. The sustained expression of a gene known as an embryonic transcription factor in adult tissue initially seems counterintuitive. However, *TBX4* has been implicated as a transcription factor of importance in human adult lung fibroblasts.^58^ Furthermore, *TBX4* has been shown to have decreased expression in human lung cancer associated fibroblasts in comparison to matched normal lung fibroblasts^58^, providing preliminary support that *TBX4* expression may be involved in FP biology.

In comparison to previous work conducted by Duffy et al.^32^ we see a strong correlation with the gene expression differences identified in the current study, despite geographical and methodological differences. Duffy et al. previously implicated the Wnt signaling pathway as a potential therapeutic avenue for FP and our current findings reiterate this potential. While we did not replicate the finding of Duffy et al. of an upregulation of WNT5A in FP tumors, among the top differentially expressed genes in this study several are involved in Wnt signaling pathways. This includes upregulated genes *CTHRC1*, *NLRC5* and downregulated *LGR5*. *CTHRC1* has been shown to selectively activate the WNT/planar cell polarity pathway^59^ and that this may have a role in cervical cancer^60^. *NLRC5* has been identified as a regulator of the Wnt/β-catenin signaling pathway in clear cell renal carcinoma^61^, and in the current study we observe increased expression of *NLRC5* in FP (log2FoldChange = 3.39, padj = 3.71×10^−12^). Upregulation of *NLRC5* in FP may represent a physiological anti-tumor response in sea turtles. Studies have shown that *NLRC5* is a transcriptional regulator of MHC class I genes^62^ and through this, activates cytotoxic CD8^+^ T cells as part of an anti-tumor immune response^63^. The increased tumor expression of *NLRC5* is associated with better outcomes in human cancer^64^ and is a target of interest in anti-tumor immunity in transmissible tumors of another endangered species; the Tasmanian devil^65^. Furthermore, downregulation of *LGR5* in FP (log2FoldChange = −4.56, padj = 1.03×10^−9^) may also demonstrate evidence of an anti-tumor response through regulation of the Wnt/β-catenin signaling pathway, as increased expression of *LGR5* is associated with poor prognosis in glioma^66^, breast cancer^67^ and oral squamous cell carcinoma.^68^ The consistent identification of the alteration of genes involved in Wnt signaling in FP indicates that further in-depth exploration of this pathway in FP is warranted.

Moving beyond individual genes, we next took a systems biology based approach drawing upon the human centered knowledge bases of Gene Ontology (GO) and the Kyoto Encyclopedia of Genes and Genomes (KEGG) pathways. We annotated all differentially expressed genes where possible with their closest orthologous human protein match to enable a comprehensive analysis within the human centered knowledge bases. The GO based enrichment analysis identified 465 nominally significant ontology terms within the gene set. These were primarily immune system based, and among the top 20 terms when considering the genes overlapping between them two functional modules emerged. One module involved the most significantly enriched GO biological process (BP), GO:0009617, response to bacterium (padj = 1.77×10^−13^). While the other module was comprised of immune related processes. Enrichment of genes involved in biological responses to bacterial infection is intuitive given that FP tumors, as an externally exposed and compromised tissue have potential to become infected with environmental bacteria. This bacterial infection interplays with tumor-based processes meaning that the enrichment of immune related GO terms is likely driven by tumor biology but also contributed to by bacterial infection. This is also reflected in the KEGG pathway-based analysis where the GSEA identified that the most significantly enriched pathway was hsa04060, cytokine-cytokine receptor interaction (padj = 8.72×10^−7^). It is well established in humans that cytokines have a fundamental role in the tumor microenvironment as well as inflammation and infection control.^69^ Here in FP, it is likely that the contribution of cytokine pathways is again an interplay between immune processes related to bacterial infection, and those related to tumorgenicity. This is supported by histopathological examination of the tumors in this study showing various degrees of lymphocytic infiltration. Given these findings, a fundamental question arises: why is this immune response insufficient to result in tumor regression in individuals that exhibit progressive disease?

In relation to this question, given that the differentially expressed genes at both the gene level and the systems level implicate significant involvement of the immune system in FP tumors, we explored immune checkpoint molecules in this data as a potential future therapeutic avenue for FP treatment. The inhibition of immune checkpoint molecules as a therapeutic approach for the treatment of cancer is a rapidly growing field.^70^ We targeted a non-exhaustive set of 13 established and emerging immune checkpoint molecules identified from the literature for assessment in the current dataset. Gene expression of three checkpoint molecules, BTLA, PD-L2 and LAG3 was significantly upregulated in FP tumors. Each of these molecules represents a potential drug target opportunity to explore in the treatment of FP. Trialing an existing human based inhibitor that is injectable directly into FP tumors may substantially improve the burden of FP management in sea turtle rehabilitation but would require significant investment and carefully designed clinical trials to identify a chemotherapeutic regimen. Given the recent increases in stranded sea turtles with FP requiring care in some regions, a therapy that facilitates rapid rehabilitation and return to the environment could be very beneficial.

Here we have described the transcriptomic profiling of FP tumors in green sea turtles, building upon prior work from Duffy et al.^32^ but also identifying novel insights into this disease. While in comparison to human tumors much remains unknown about the basic biology of sea turtle FP there is significant opportunity to further apply alternative approaches previously established in humans and model organisms to further our understanding of this disease. Research approaches involving single cell sequencing, long read sequencing, metagenomics and proteomics all stand to substantially benefit the FP knowledge base. Investment in this area, may even offer returns to human oncology.^71^ Wildlife health problems, such as FP, are increasing and will require a multidisciplinary approach to their mitigation.

## METHODS

### Sample collection

Sampling of green sea turtles (*Chelonia mydas*) was carried out under permit number TE181762-4 from the U.S. Fish and Wildlife Service and with the ethical approval of the University of Texas Rio Grande Valley’s Institutional Animal Care and Use Committee (IACUC). The 9 turtles involved in this study were undergoing rehabilitative care at Sea Turtle Inc., South Padre Island. Three turtles, at the time of collection had external FP tumors and tumor burden was scaled according to the National Oceanic and Atmospheric Administration (NOAA) Ordinal Scale for Grading Fibropapillomatosis by Photographic Comparison as described in Stacy et al. 2018^34^. From these three turtles, external FP tumors were sampled after removal during rehabilitative surgery using CO_2_ laser resection. From all turtles a single healthy control skin sample was collected using 3-6mm punch biopsies of the hind flipper. Tissue samples were collected into RNA-later (Qiagen) and stored at −80°C until processing. All turtles sampled in this study were juvenile green turtles of unknown sex. Table 1 summarizes the individual characteristics of turtles in this study.

### Histopathological studies of tissue samples

For each tumor sample collected, approximately 50% of the sample was fixed in a 10% formalin solution. Tissues were processed into paraffin blocks, sectioned by 5 μm, mounted onto glass slides and stained with hematoxylin and eosin using routine methods. Diagnosis of fibropapilloma was confirmed based on previously described histopathological features.^72^

### RNA extraction and RNA-seq

Tumor and healthy tissue samples were homogenized using a rotor-stator homogenizer and total RNA was extracted using the Qiagen AllPrep DNA/RNA Mini kit, according to the manufacturer’s instructions. RNA quality was assessed on a 2200 TapeStation System (Agilent Technologies) with sample RIN values ranging from 6.4 to 9.6 (μ=8.4). mRNA sequencing libraries were generated from up to 250ng of total RNA and were prepared using a KAPA RNA HyperPrep kit (Roche Diagnostics) with polyA selection and indexed using a KAPA Dual-Indexed adapter kit (Roche Diagnostics), according to manufacturer’s instructions. Prepared libraries were evaluated via the 2200 TapeStation System and were pooled for paired-end 100bp sequencing on a HiSeq 2500 system (Illumina).

### Quality control of RNA sequence data

Sequence data was demultiplexed using bcl2fastq. Raw sequencing reads were processed with Trim Galore (version 0.6.5, https://www.bioinformatics.babraham.ac.uk/projects/trim_galore/), using Cutadapt^73^ to remove adapter sequences and indexes from reads and to exclude low quality sequences using a Phred score of 30 and discarding reads with lengths shorter than 25bp, unpaired reads were not retained. FastQC (version 0.11.9, (https://www.bioinformatics.babraham.ac.uk/projects/fastqc/) was used to assess the data quality of the processed reads. To avoid potential bias in analysis two individual samples were excluded due to overall differences in total sequencing content, one tumor tissue (FP03-05, Beach Bum) had over 83 million more paired-end reads than the next highest sample and one healthy tissue (FP08-01, Robocop) had over 2.5 million less paired-end reads than the next lowest sample. In the remaining 33 samples, the mean and median number of paired-end sequencing reads was 26,607,522 and 23,683,506 respectively, ranging from 4,447,485 to 70,675,924 reads.

### Sequence alignment and quantification of turtle gene expression

The CheMyd_1.0 reference assembly for *C. mydas* was obtained from NCBI [GenBank assembly accession number: GCA_000344595.1].^74^ Reads were aligned to the reference assembly using HISAT2 (version 2.2.0)^75^ with an overall alignment per sample ranging from 75.26% to 88.29% (μ=84.19%). Transcript abundance was quantified in each sample using htseq-count (version 0.11.2)^76^ at the gene level according to the defined genomic features of the CheMyd_1.0 assembly.

### Differential expression analysis

Differential expression analysis was conducted with DESeq2 (version 1.28.1)^77^. Genes with low counts were filtered from the analysis, retaining genes with more than 5 counts in 2 or more samples. This corresponded to 17,698 genes (83%). To calculate factors of unwanted variation in this sample using RUVSeq (version 1.22.0)^78^, a first pass differential expression analysis was conducted to identify a set of *in-silico* empirical negative control genes. This corresponded to a model comparing 25 tumor samples to 8 healthy tissue samples with inclusion of a turtle covariate in the model to account for multiple tumor samples obtained from individual sea turtles. Using a test statistic p-value of > 0.5 to identify genes not differentially expressed in the comparisons there were 5,730 genes identified as empirical control genes from the analysis. Using RUVSeq two factors of unwanted variation were calculated from the data. These factors were then incorporated into the DESeq2 analysis design and an analysis was conducted to identify differentially expressed genes with a log2 fold change threshold of 1 and a false discovery rate threshold of 0.05. Relative log expression plots of the data unadjusted and after normalization, including the two factors of unwanted variations from RUVSeq are shown in supplementary figures 1 and 2 respectively. The differential expression analysis was visualized as a volcano plot using the ‘EnhancedVolcano’ package (version 1.6.0)^79^. Correlations of previously published differential gene expression changes with the results from this study were calculated using a Pearson correlation test in R (version 4.0.0).

### Human ortholog identification

To enable pathway based and gene set enrichment analyses in this data sea turtle genes were matched to their closest matching human gene orthologs based on protein amino acid sequence homology. To accurately draw upon human biological understandings for inference in sea turtle fibropapilloma the defined set of 28,672 proteins for the CheMyd_1.0 reference was compared to the human landmark protein database using NCBI’s protein BLAST^80–82^ to identify the closest human orthologous match. Initially the blastp query was run with the following parameters: -max_target_seqs 1 -max_hsps 1 -evalue 1e-6. This identified a match for 27,961 sea turtle proteins (97.5%). For the remaining 711 sea turtle proteins, the blastp analysis was repeated reducing the threshold of the expect value (evalue) to 0.01. This matched a further 165 human proteins to turtle proteins. Finally, the evalue threshold was reduced to 1, matching a further 182 proteins. In total 28,308 protein sequences (98.7%) from the CheMyd_1.0 reference were successfully matched with a human landmark database protein.

Turtle and human protein accession numbers were converted to gene symbols using bioDBnet.^83^ A subset of 211 turtle protein isoforms of 101 individual turtle genes matched to multiple human genes. When there was concordance between the human and turtle gene symbols for one or more of the turtle protein isoforms that gene symbol was used as the match between turtle and human. For the remaining genes, the gene symbol corresponding to the blastp result with the lowest evalue was used. For evalue ties the gene symbol of the result corresponding to the longest turtle protein isoform was used.

For the remaining 364 turtle proteins that could not be matched to a human protein, the sea turtle genome gene annotation was used. For 2,233 turtle genes, including predicted tRNA molecules and other non-coding genes, a corresponding protein sequence was not available and for these genes the sea turtle genome gene annotation was retained.

Supplementary table 4 provides a mapping from CheMyd_1.0 gff_file gene identifiers through to the human based gene matches used in this project.

### Gene set enrichment analysis

Gene set enrichment analysis (GSEA) was conducted using clusterProfiler (version 3.16.1).^84^ Human gene symbols from the orthologous matching analysis of sea turtle genes were annotated with Gene Ontology (GO) terms and Kyoto Encyclopedia of Genes and Genomes (KEGG) pathways. An adjusted p-value of < 0.1 was used to identify input genes for this analysis. For each duplicate ortholog the most significant differentially expressed gene value was used in the GSEA analyses. The background gene set used was all orthologous human genes identified as matching a sea turtle gene. Test statistics were adjusted for multiple testing using the Benjamini-Hochberg method.

### Detection of chelonid alphaherpesvirus 5 sequences

The chelonid alphaherpesvirus 5 (ChHV5) reference assembly was obtained from NCBI [GenBank assembly accession number: GCA_008792165.1].^85^ Reads were aligned to the reference assembly using HISAT2 (version 2.2.0)^75^ and transcript abundance was quantified in each sample using htseq-count (version 0.11.2)^76^ at the gene level according to the defined genomic features of the ChHV5 assembly. Differential gene expression was not conducted for ChHV5 with results only reported as a presence or absence of viral transcripts in each sample.

### Data availability

The RNA-Seq data, including raw fastq reads, are deposited in NCBI (https://www.ncbi.nlm.nih.gov/) under BioProject ID: PRJNA672698 (https://www.ncbi.nlm.nih.gov/bioproject/PRJNA672698) and will be made publicly available upon publication.

## Supplementary figures

**Supplementary Figure 1.**
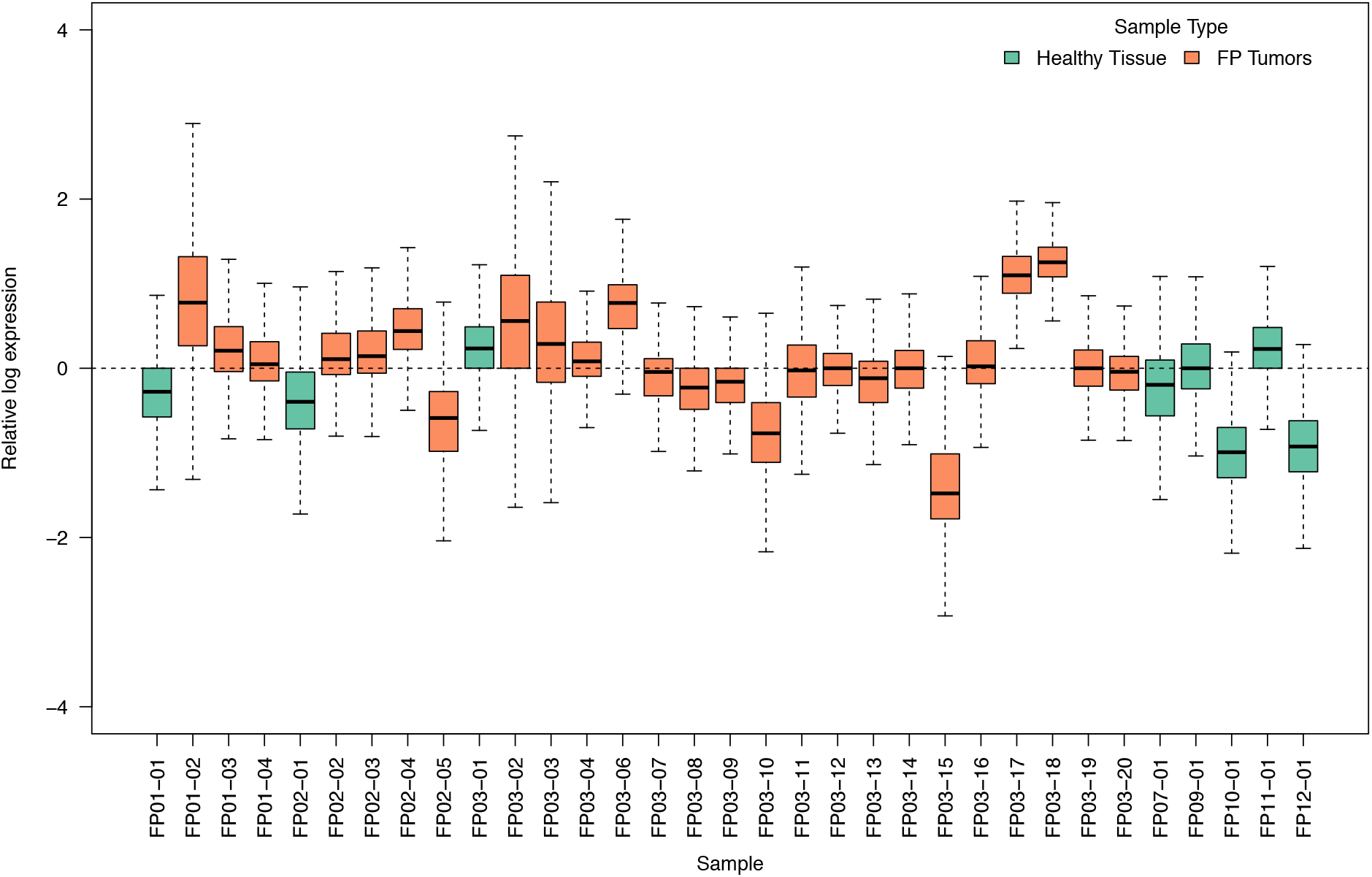
Relative log expression plot of the gene expression data per sample prior to adjustment for two factors of unwanted variation calculated using RUVSeq.

**Supplementary Figure 2.**
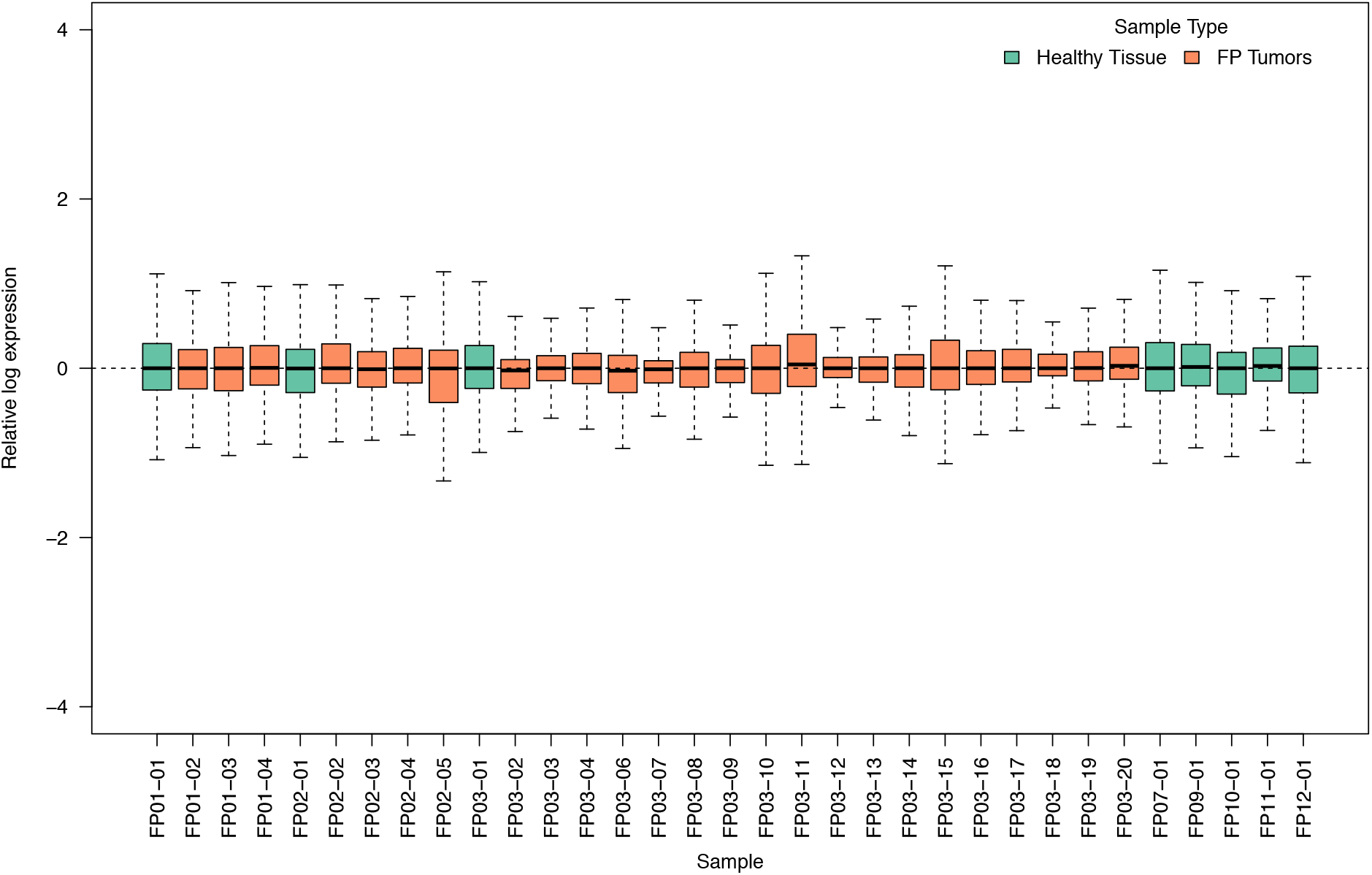
Relative log expression plot of the gene expression data per sample after adjustment for two factors of unwanted variation calculated using RUVSeq.

## ACKNOWLEDGMENTS

This work was funded by the University of Texas Rio Grande Valley Transforming Our World Strategic Plan. This work was conducted in part in facilities constructed under the support of NIH grant C06 RR020547. The co-authors wish to acknowledge the ongoing and generous contributions from supporters of Sea Turtle Inc., a non-profit sea turtle conservation, rehabilitation and education facility located on South Padre Island, Texas. Without public support for the Sea Turtle Inc. facilities, employees and associates the findings presented here would ultimately not have been possible.

## FINANCIAL AND NON-FINANCIAL COMPETING INTERESTS STATEMENT

The authors declare no competing interests. Funding agencies had no role in the conceptualization, design, data collection, analysis, decision to publish or preparation of the manuscript.

## Notes

### Competing Interest Statement

The authors have declared no competing interest.

